# Phase separation on cell surface: a mechanism of basic fibroblast growth factor signal transduction with heparan sulphate

**DOI:** 10.1101/2021.05.24.445073

**Authors:** Song Xue, Fan Zhou, Tian Zhao, Huimin Zhao, Xuewei Wang, Long Chen, Jin-ping Li, Tianwei Tan, Shi-Zhong Luo

## Abstract

Liquid-liquid phase separation (LLPS) driven by weak, multivalent interactions among biomolecules is an important means of cellular compartmentation and plays a central role in cellular processes including stress resistance, RNA processing and other cellular activities. Coordination of the condensates and inner membrane was recently revealed, mediating intracellular processes like cell signalling and cargo trafficking. Intracellular LLPS has been observed extensively *in vivo*, whereas LLPS in extracellular compartments has not been reported under physiological conditions. Here we show, for the first time, that basic fibroblast growth factor (bFGF) undergoes LLPS on the cell surface by interacting with heparan sulphate proteoglycans (HSPG) and the phase transition is required for effective downstream signalling. The condensation is driven by multivalent interactions between bFGF and sulpho-groups on heparan sulphate (HS), and dimerization and oligomerization of bFGF promote the LLPS process. Compared with free bFGF, phase separated bFGF with HS showed higher thermo stability, providing a potential mechanism for the preservation of bFGF activity. Furthermore, we have found that downstream signalling is triggered by phase separation of a ternary complex formed by bFGF, HSPGs and FGFR on cell surface. Our results revealed a molecular mechanism that HS can serve as a platform to promote extracellular proteins like bFGF to condensate on outer membrane, consequently coordinating the signal transduction activities. This novel finding expands the horizons of phase separation *in vivo*, providing a new dimension on how HSPG may regulate extracellular protein behaviour and cell signalling.

## Introduction

LLPS plays critical roles in cellular processes, including formation of membrane-less organelles, construction of stress granules, regulation of genome organization and control of synaptic signaling^1, 2^. LLPS driven by weak multivalent interactions promotes formation of distinct functional condensates inside cells^2^. Those condensates can be assembled in the cytoplasm and nucleoplasm, such as processing bodies involved in RNA turnover, stress granules reacting against harmful conditions, and chromatin organization^1, 3^. LLPS can also occur on plasma membrane or endoplasmic reticulum, which provides platforms for condensates formation, facilitating cell signalling, tight junction formation and synaptic transmission^4, 5, 6, 7, 8^. Most of current reported LLPS *in vivo* happens in the intracellular environment, which might due to the high protein density in cells and other biomolecules is amenable to form multivalent weak interactions that drives phase separation^1^. As the mechanisms that promote and regulate LLPS *in vitro* and inside cells are getting elucidated^9, 10^, it remains an open question whether LLPS can happen in extracellular environments. Here, we demonstrate that basic fibroblast growth factor (bFGF) undergoes LLPS, which is critical for its signalling activity.

The fibroblast growth factors (FGFs) are a family of proteins with key roles in variety of processes, such as embryonic development, tissue regeneration and wound healing^11, 12^. The member of FGF2, also named basic FGF (bFGF) from its rich basic residuals, is an important regulator of cell growth and differentiation under physiological and pathological conditions^11, 12^. In extracellular matrix (ECM), bFGF bounds with heparan sulphate proteoglycans (HSPGs) for storage and is released to cell surface by matrix degradation, serving as a mechanism in response to injury or tissue reorganization^13^. bFGF can also be secreted by adipocytes, and then diffuses to nearby cells. Secreted and released bFGF diffuses to nearby cells and is sequestered by HSPG that are tethered to the cell surface. On responsive cells, bFGF forms a ternary complex by binding to HSPG and fibroblast growth factor receptor (FGFR), and triggers downstream response^13^. It also has been proposed that HSPG can associate with bFGF with a low affinity in a dynamic manner to assist receptor binding in close proximity^14^. Our study revealed that the weak binding promotes bFGF to form a condensate with HSPG via phase separation, and this condensate can further incorporate FGFR to form the active receptor complex.

## Results and Discussion

### Heparin promotes bFGF to undergo liquid-liquid phase separation

Proteins capable of phase separation usually contain intrinsically disordered regions or tandem repeats. bFGF contains disordered N- and C-terminal sequences^15^ and can assemble into oligomers^16^, which may provide sufficient multivalent weak interactions for LLPS. To explore the possibility of phase separation, enhanced green fluorescent protein fused bFGF (eGFP-bFGF) was expressed from *E. coli* and purified (Extended Data Fig. 1a). This eGFP fused bFGF has similar activity to stimulate cell proliferation as the wild-type protein (Extended Data Fig. 1b). The status of purified protein at various concentrations of PEG-8000 was examined under confocal microscope. Protein droplets formed and enlarged as the concentration of PEG increase to 10%, indicating LLPS. Further increase of PEG concentration led to a decrease in both the size and quantity of the droplets and an increase in irregular aggregation (Fig. 1a). The solution turbidity also peaked at 10% PEG concentration and dropped at lower or higher PEG concentration, which is consistent with the image-based analysis (Fig. 1b). The droplets exhibited typical liquid property. In fluorescence recovery after photobleaching (FRAP), fluorescence of the droplets quickly recovered, suggesting that their contents have high fluidity (Fig. 1c). A subset of droplets also merged during observation, consistent with their liquid-like nature. In contrast, eGFP-bFGF formed solid aggregation in 20% PEG, distinguished by the irregular morphology and irreversible FRAP results (Fig. 1d). The results demonstrated that bFGF tends to phase separate in a mild crowded environment but form aggregates if the environment is overcrowded. As the extracellular environment is generally less crowded than the cytosol and bFGF is present at a low concentration, the finding suggests that bFGF can phase separate under physiological conditions.

**Fig. 1.**
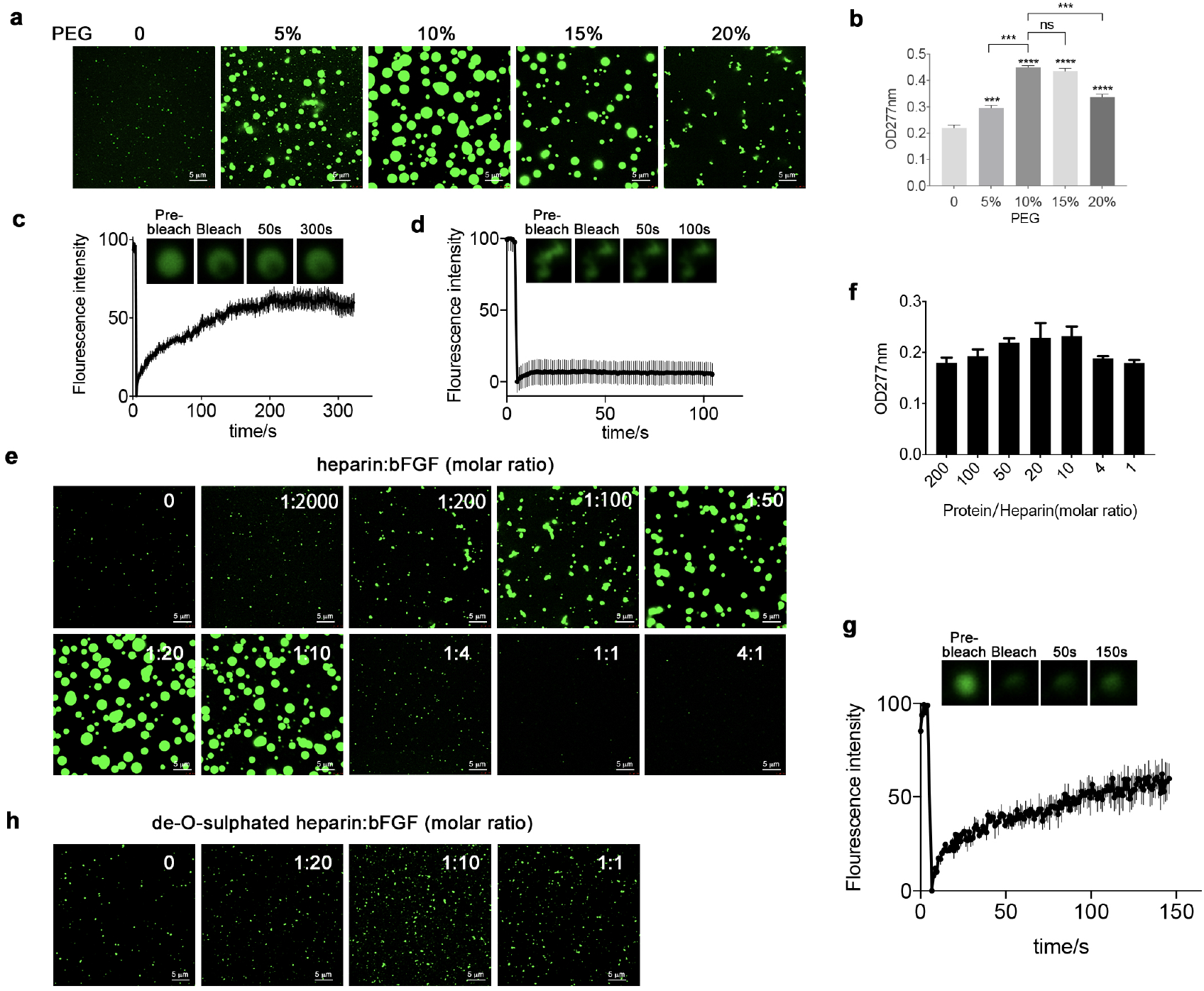
bFGF undergoes phase separation in the presence of PEG or heparin. **a**, Confocal microscopy images of assembling status of 5 uM eGFP-bFGF with 0-20% PEG. Scale bar=5 µm. **b**, Turbidity measurement of eGFP-bFGF with 0-20% PEG. **c**, FRAP of the droplets formed by eGFP-bFGF with 10% PEG. ns, not significant; *p < 0.05, **p < 0.01, ***p < 0.001, and ****p < 0.0001. Comparisons among groups were performed using one-way ANOVA with Krusk–Wallis test or unpaired two-tailed Student’s t-tests with GraphPad Prism 7.0. **d**, FRAP of the aggregates formed by eGFP-bFGF with 20% PEG. **e**, Confocal microscopy images of assembling status of 5 uM eGFP-bFGF mixed with heparin at different ratios. **f**, Turbidity measurement of eGFP-bFGF with heparin. **g**, FRAP results of the droplets formed by heparin:eGFP-bFGF = 1:20. **h**, Confocal microscopy images of assembling status of 5 uM eGFP-bFGF mixed with de-O-sulphated heparin at different ratios.

Though PEG and other crowding agents are commonly used to mimic crowded intracellular environments^17^, bFGF is extracellular and the environment is distinct. HSPGs are abundant in the ECM and cell surface and are known to interact with bFGF.^18^ As the negatively charged liner polysaccharide can provide multiple bFGF binding sites and result in more multivalent interactions, we hypothesize that interaction between bFGF and HSPG may promote LLPS. To test this hypothesis, we mixed heparin, a commonly used HS mimetic, with 5 µM eGFP-bFGF at different ratios and examined the behavior of the mixture. As the molar ratio of heparin to eGFP-bFGF increased to 1:10, more and lager droplets were formed, indicating stronger phase separation (Fig. 1e), suggesting that heparin promotes the phase separation of eGFP-bFGF. Higher concentrations of heparin prevented droplets formation, demonstrating a “reentrant” behaviour, similar to the effects of RNA in the phase separation of many RNA binding proteins^19, 20^. Indeed, heparin or HS share similar structural properties with RNA, such as rich negative charges and linear repetitive structure, and may promote LLPS following a similar mechanism as RNA. Change in turbidity is also consistent with the observations under microscopy (Fig. 1f). These observations suggest that heparin promotes bFGF phase separation. It has been reported that binding to heparin can increase the stability of bFGF by protecting it from proteolytic degradation^21, 22^. We further tested whether heparin-induced phase separation can increase bFGF thermostability, as bFGF tends to unfold quickly at body temperature. eGFP-bFGF was incubated under different temperatures with or without heparin. With increased temperature, bFGF alone formed aggregates, while the samples with heparin remained in liquid phase at 42 °C (Extended Fig. 2). The results suggest that heparin-induced phase separation increased thermostability of bFGF.

**Fig. 2.**
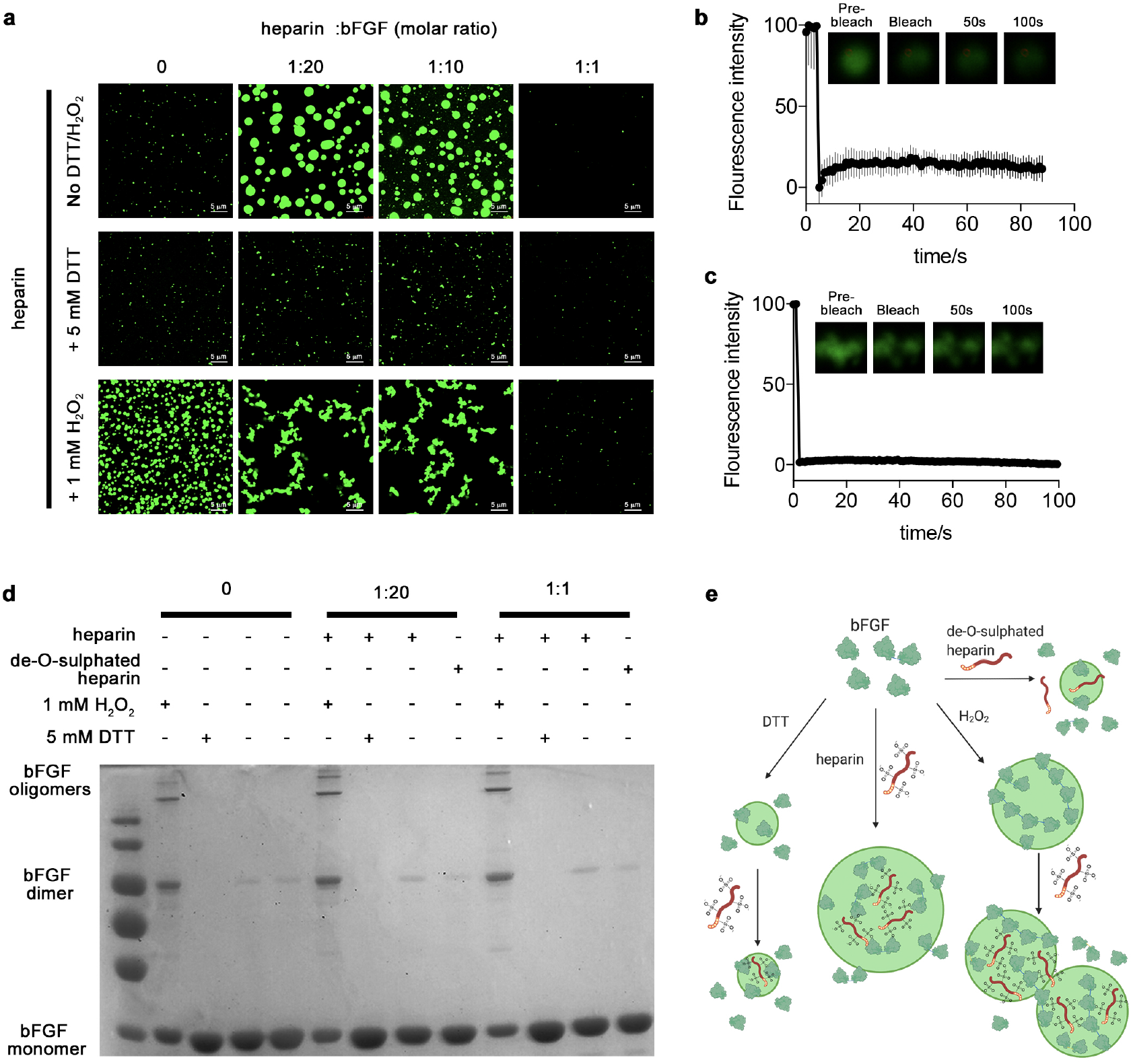
Interaction of bFGF and heparin. **a**, Confocal microscopy images of bFGF condensates with various ratios of heparin with 5mM DTT or 1 mM H_2_O_2_. Scale bar=5 µm. **b**, FRAP of the condensates formed by eGFP-bFGF oxidized in the presence of 1 mM peroxide. **c**, FRAP of the condensates formed by oxidized eGFP-bFGF : heparin =1:20. **d**, Non-reduced SDS-PAGE of the bFGF with different oxidative conditions and heparin concentrations, showing its assembling status. **e**, Schematic illustration of the proposed mechanism of bFGF assembling under different oxidative status with heparin/ de-O-sulphated heparin.

Natural heparin is highly sulphated and the charged sulpho group is likely critical for the interaction with bFGF. It has been reported that 6-O-sulfate groups is critical for HS to promote the activity of bFGF.^23^ To assess the importance of sulfation in heparin, we used a de-O-sulphated heparin (OD-heparin)^24^. Addition of the OD-heparin failed to induce LLPS of bFGF (Fig. 1h). This indicates that the phase separation of bFGF is mainly driven by the multivalent interactions between the protein and highly negatively charged sulpho group on heparin.

### Oxidation status of bFGF affects its LLPS

Before exploring the behaviour of bFGF at the cellular level, we investigated the structural mechanisms of bFGF-heparin interaction. bFGF has two exposed cysteines and can exist as dimers and oligomers. The two exposed thiol groups form disulphide bonds under oxidizing conditions and become unstable with aging of the protein^25^. Dimerization is important to bFGF activity, which further induces FGFR dimerization and activation^16^. Chemically conjugated multivalent bFGF was also reported to be more potent to stimulate cell proliferation since it is believed to bring more receptors in close proximity, assisting the dimerization of FGFR for signalling transduction^26^. To examine whether reduction and oxidation of the cysteine residues in bFGF affect its phase separation, we added 1 mM hydrogen peroxide or 5 mM 1,4-Dithiothreitol (DTT) to the eGFP-bFGF solutions with various concentrations of heparin (Fig. 2a). Notably, phase separation of bFGF was essentially inhibited by DTT regardless with or without heparin. With hydrogen peroxide added, the bFGF alone formed round condensates (Fig. 2a). However, FRAP assay showed no mobility of those condensates, indicating that they were gel-like entities crosslinked by disulphide bonds (Fig. 2b). The round shapes of the condensates implied that phase separation may occur in early stage and the droplets solidified quickly. Addition of heparin promoted the formation of clusters of condensates having no mobility either (Fig. 2c). This indicates that the oxidative condition resulted in larger population of dimers or even oligomers as confirmed by SDS-PAGE (Fig. 2d), which possesses more multivalent interactions of bFGF and heparin, favourable for aggregates formation. Endogenous bFGF exists mainly in monomers, with a small portion of dimers and multimers. We have reasons to believe that the population of different assembling status of bFGF in physiological conditions can provides sufficient multivalency for phase separation but not for aggregation as illustrated in Fig. 2e.

### bFGF phase separates on cell surface mediated by heparan sulphate

As bFGF can undergo phase separation in presence of heparin *in vitro*, we further tested phase separation of bFGF on cell surface, where heparan sulphate is almost ubiquitous. We applied 500 nM eGFP-bFGF to several cell lines, including mouse neuroblastoma cells (N2a), Chinese hamster ovary cells (CHO-K1) and mouse embryonic fibroblast cells (MEF, BALB/C-3T3) cells to examine phase separation by confocal microscope. We found bright condensates formed on the edges of the cells incubated with eGFP-bFGF. Z-stack scanning further confirmed existence of the condensates on the cell surface (Fig. 3a). As control, eGFP alone was applied to the cells and no droplet formation was observed (Extended Fig. 3). FRAP experiment showed good fluorescence recovery, indicating liquid-like properties of the condensates (Fig. 3b and Extended Fig. 3b). Larger number of droplets were observed on MEF cells compared with the other two strains, since HS was more abundant on MEF, indicating that HS induced LLPS. Moreover, in cells stained with anti-heparan sulphate antibody, eGFP-bFGF condensates overlapped significantly with heparan sulphate, suggesting that heparan sulfate likely mediated the formation of bFGF condensates (Fig. 3c). Further, we treated MEF cells with heparinase III to digest heparan sulphate on cell surface before adding eGFP-bFGF, and saw a significant decrease of the condensate amount (Fig. 3d). Moreover, in CHO-677, a cell line lacking heparan sulfate^27^, no condensate was detected, in contrast to the clear condensates observed on the CHO-K1 WT cells (Fig. 3e). These observations suggest that heparan sulphate is required for phase separation of bFGF on the cell surface. Phase separation increases effective concentration of bFGF within the phase and may promote its interaction with the receptor and affect downstream signalling.

**Fig. 3.**
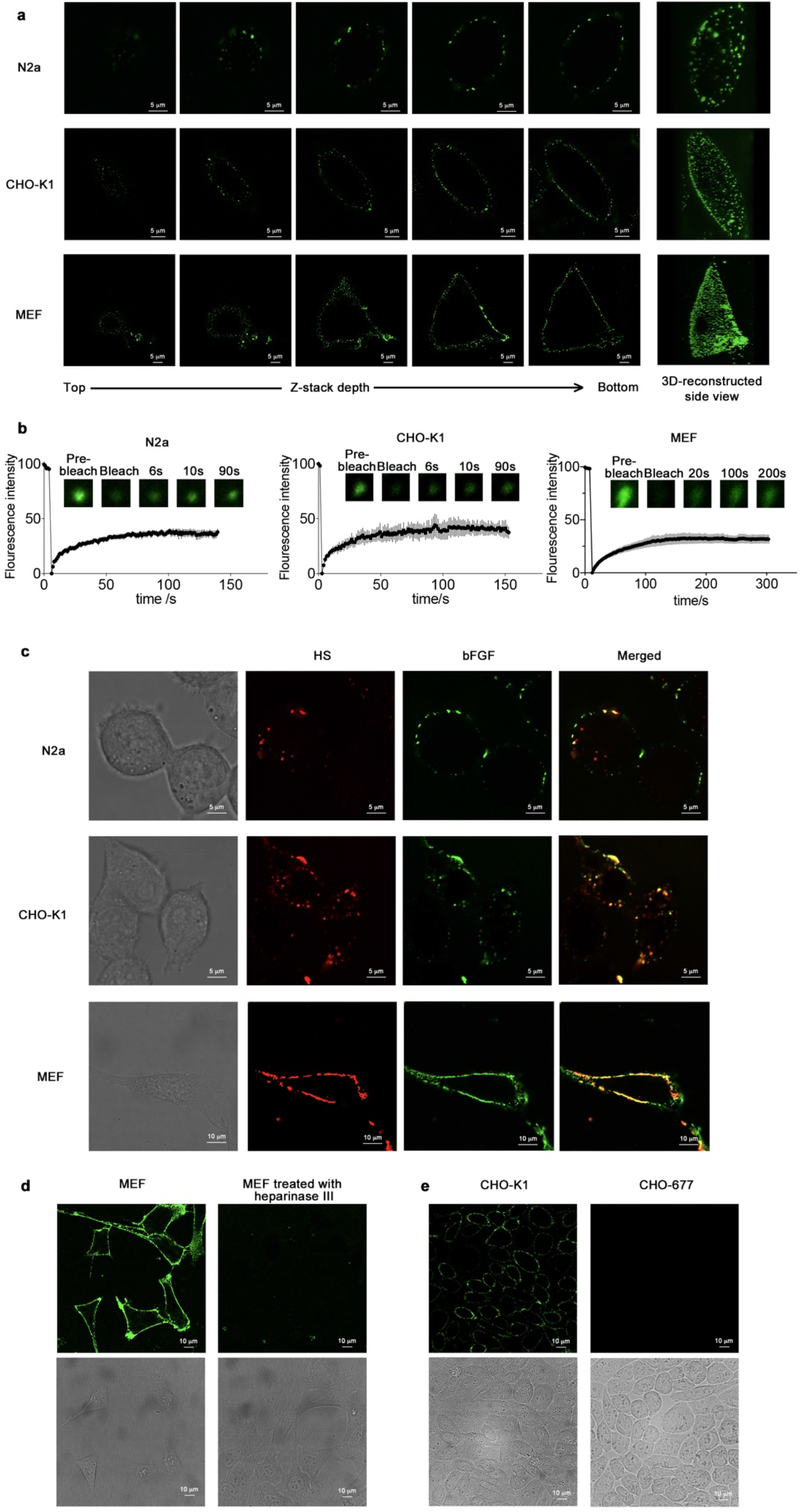
bFGF phase separates on cell surface with heparan sulphate. **a**, Z-stack scanning of droplets formation on the cell surface. N2a, CHO-K1 or MEF cells incubated with eGFP-bFGF were imaged under confocal microscope with the Z-stack method, showing droplets condensation on the cell surface. Scale bar=5 µm. **b**, FRAP of the condensates formed by eGFP-bFGF. **c**, Confocal microscopy images of heparan sulphate (HS, red) immunostaining and eGFP-bFGF (green) on N2a, CHO-K1 and MEF cells. Scale bar=5 µm for N2a and CHO-K1 cells; 10 µm for MEF cells. **d**, Confocal microscopy images of eGFP-bFGF phase separation on MEF cell surface with or without treatment with heparinase III. Scale bar=10 µm. **e**, Confocal microscopy images of eGFP-bFGF phase separation on the surface of wildtype CHO-K1 and the HS deficient CHO-677 cells. Scale bar=10 µm.

### LLPS of bFGF facilitates signal transduction

FGFs exert activities through binding to their receptors, and heparan sulphate functions as a co-receptor to form HSPG-bFGF-FGFR ternary complex^28, 29^. Since bFGF phase separate with HS on the cell surface, we suspected that the complex may co-condensate. FGFR antibody was used to label the receptor and the formation of FGFR and bFGF condensates was examined under confocal microscope (Fig. 4a). FGFR co-localized with bFGF condensates. Moreover, some bFGF condensates do not have FGFR signal, indicating the low affinity binding of bFGF with HSPG for its sequestering^14, 30^. Next, we examined the impact of ternary phase separation on downstream signalling transduction. bFGF can activate multiple downstream signalling pathways, including Ras-ERK and PI 3-Kinase-Akt pathway^31^. Here we examined the phosphorylated ERK as a marker for the signalling activation. We applied different concentrations of bFGF to MEF cells, monitored its phase separation under confocal microscope and quantified p-ERK level with Western blot. To determine the effect from LLPS, we added 500 µg/mL heparin to the cells with bFGF since the phase separation of bFGF can be inhibited by high concentration of heparin *in vitro* (see Fig. 1). This inhibition was observed on cell surface as well (Fig. 4b). As expected, the ERK phosphorylation increased along with bFGF through in a dose-dependent manner (Fig. 4c and d). When phase separation was inhibited by excess heparin or heparan sulphate was digested by heparinase, ERK phosphorylation was much diminished. The results strongly suggested that phase separation is essential for activation of the downstream pathway of bFGF. Considering that bFGF exists in a low concentration (1 ng/mL) in human tissues^32^, there may be a mechanism to amplify its effects for the high efficacy. We believe that LLPS driven by the interactions between bFGF and HSPG effectively increase the local concentration of bFGF around FGFR, augmenting growth signal transduction (Fig. 4e). Additionally, our findings also provided a new model to explain the mysteries synergistic effects of heparin to bFGF, which heparin stimulates bFGF’s effect at low concentrations while reduces it at high concentrations.^33, 34^

**Fig. 4.**
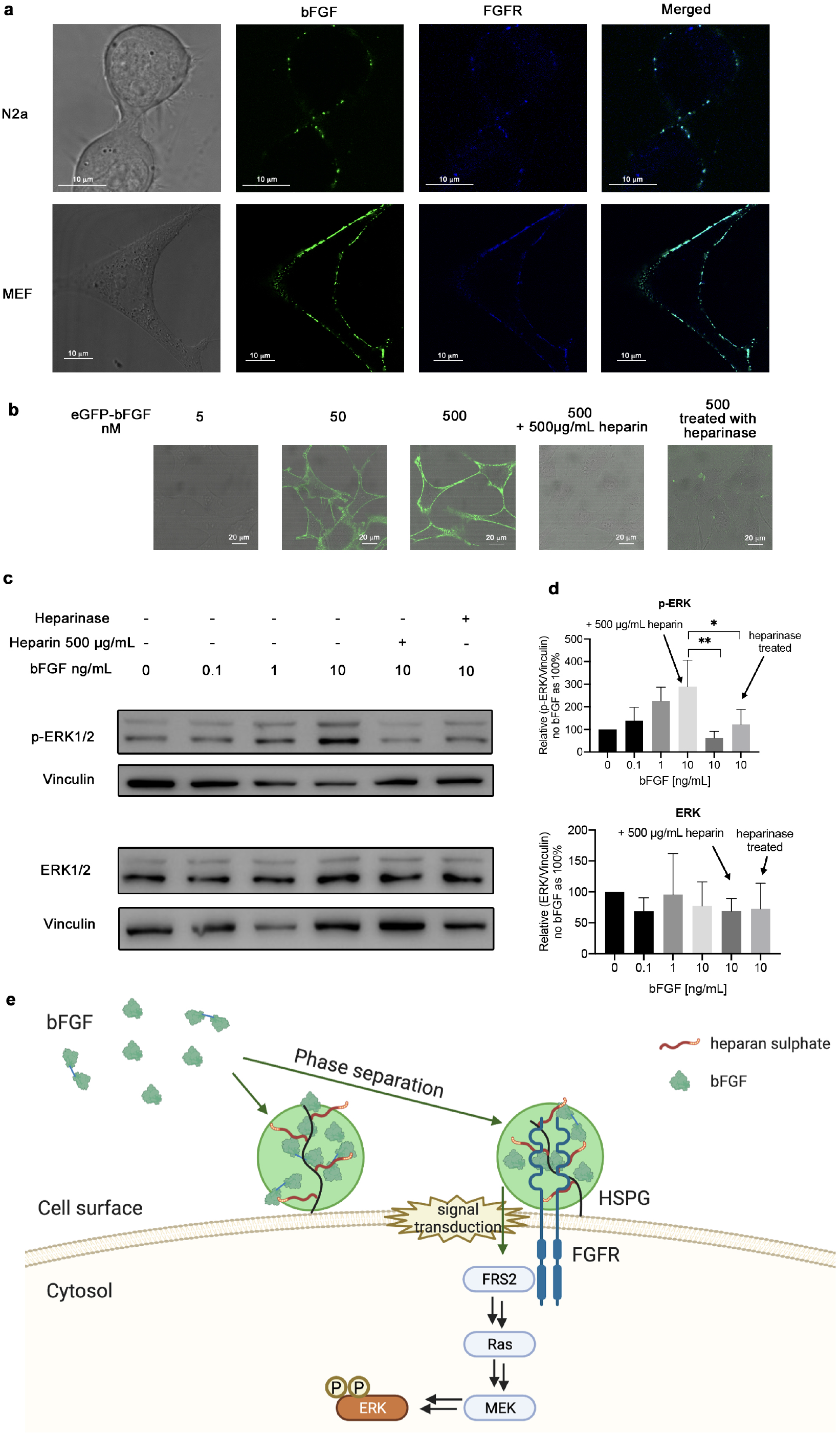
The phase separation of bFGF triggers signalling transduction. **a**, Confocal microscopy images of fluorescence-antibody FGFR (blue) and eGFP-bFGF (green) on the surface of N2a or MEF cells. Scale bar=10 µm. **b**, Microscopy images showing phase separation of different concentrations of eGFP-bFGF on MEF cells, with samples treated with 500 µg/mL heparin or heparinase. Scale bar=20 µm. **c**, Western blot showing ERK phosphorylation stimulated with bFGF at different concentrations, indicating the activation of Ras-ERK pathway. Added heparin (500 µg/mL) inhibited the downstream signalling. **d**, Quantification of ERK expression and phosphorylation with bFGF phase separation. N=4, unpaired t-test, *p < 0.12, **p < 0.033. **e**, Schematic illustration of the proposed model that bFGF phase separation along with HS on the cell surface for its capture and signalling activities.

## Conclusions

Phase separation has gained recognition in multiple cellular processes, including membrane-less organelles formation and chromatin condensation. In these processes the intracellular environment is required to provide crowding effect and promote phase separation by multivalent weak interactions. Our study, for the first time, revealed phase separation activity on cell surface, where heparan sulphate serves as a platform to induce phase separation of bFGF. In this process, bFGF is recruited and condensed into a distinct phase, which further facilitates the formation of bFGF-HSPG-FGFR ternary complex to activate downstream signal transduction as well as stabilization of bFGF. Phase separation on the cell surface thus represents a distinct mechanism for regulation of bFGF signalling.

Heparan sulphate is required for the interaction between a wide range of different cytokines and their receptors. HS mediated phase separation thus may apply to other signalling pathways and reshape the downstream response. Molecules like heparan sulphate can act as platforms to enrich signalling molecules and tune signal transduction. In addition, our findings also suggest that phase separation occurs not only in the intracellular environment but also in the extracellular environment, and has a potentially huge impact on extracellular physiology.

## Extended data figures and tables

**Extended Data Figure 1.**
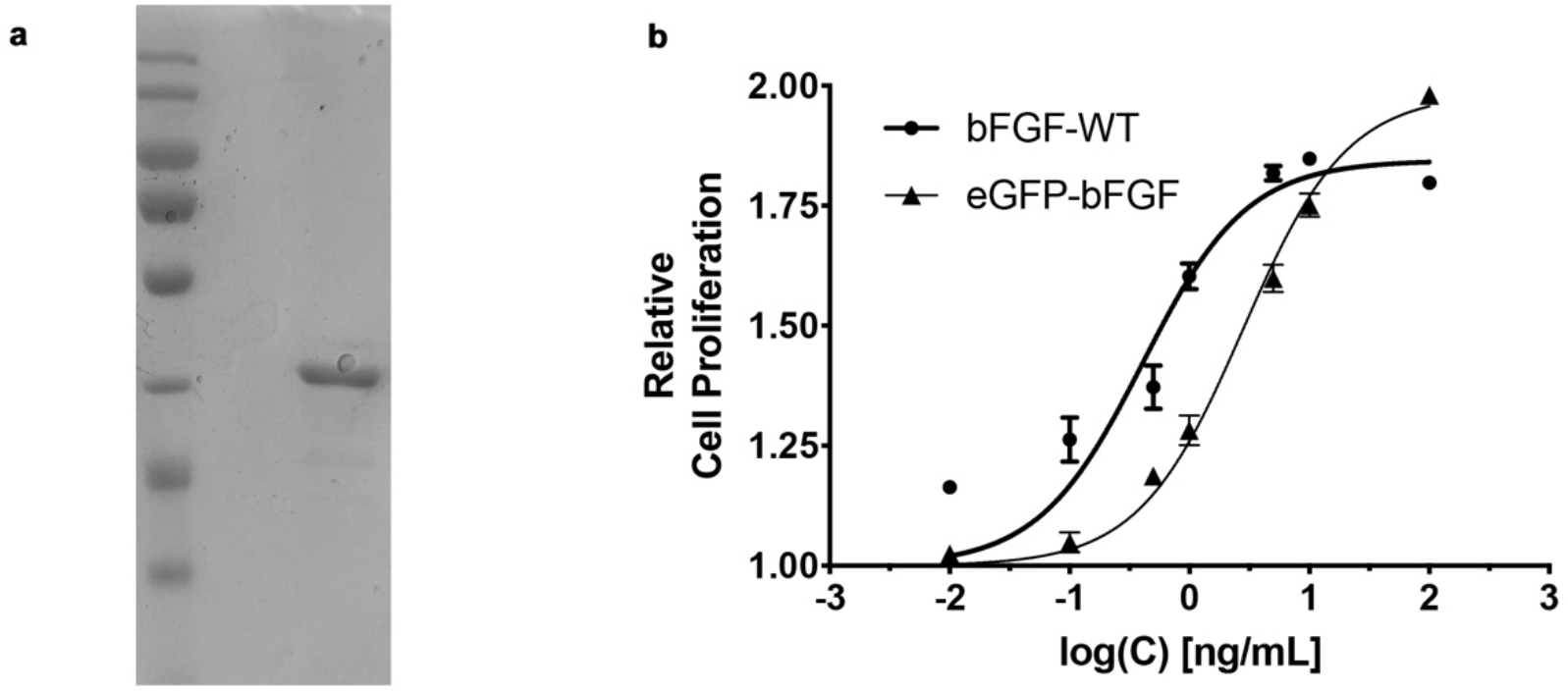
Characterization of eGFP-bFGF. **a**, SDS-PAGE showed the molecular weight and purity of eGFP-bFGF. **b**, Expressed eGFP-bFGF has similar activity as commercially tag-free bFGF, N=3.

**Extended Data Figure 2.**
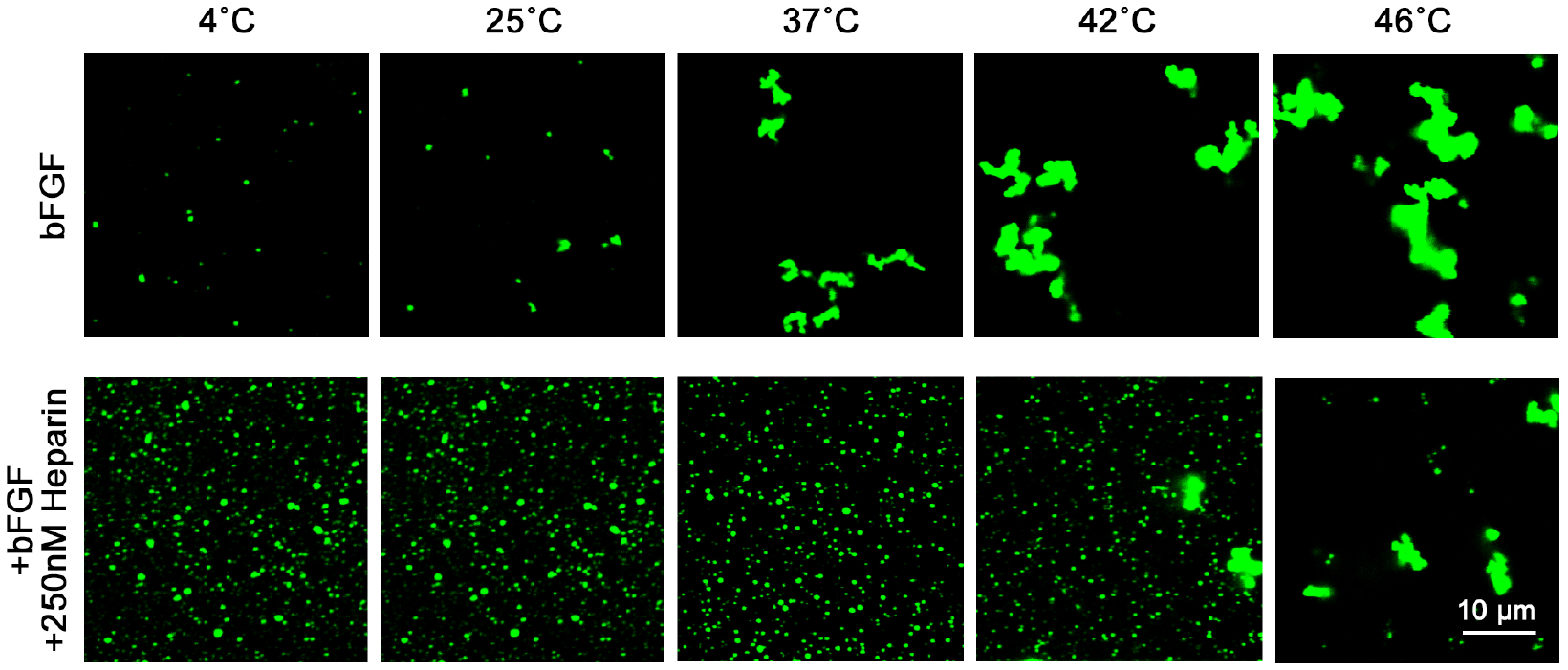
Phase separation of bFGF with heparin increases the thermo stability of bFGF. Aggregates formed in the solution of 5 µM of bFGF as the temperature increased from 4 to 46 °C. With 250 nM of heparin added, bFGF existed in a phase separation status at the temperature from 4 to 42 °C and aggregates formed at 46 °C. Scale bar=10 µm.

**Extended Data Figure 3.**
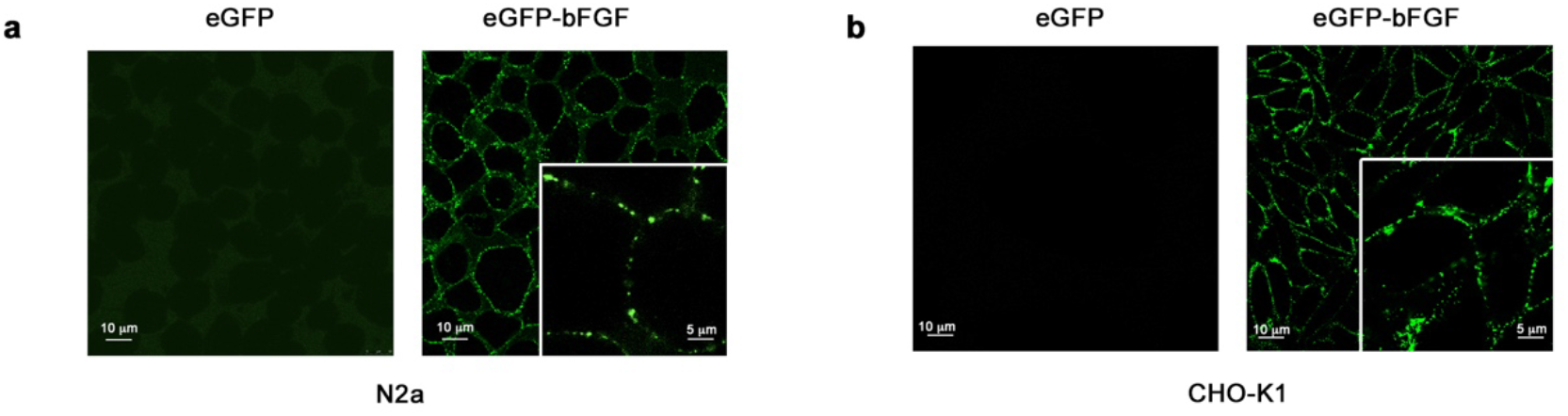
Confocal microscopy images of bFGF phase separation on cell surface, with eGFP as control. **a**, N2a cells; **b**, CHO-K1 cells. Scale bar=10 µm (5 µm for the zoomed part).

## Methods

### Protein expression and purification

His-eGFP or His-eGFP-bFGF gene were constructed in pET-28a vector (Miao Ling Plasmid, China) and transformed into in *E*.*coli* (Transetta (DE3), TransGen Biotech, China) cells for expression. Cells were then grown to optical density of 0.9-1.2 at 37 °C and induced with 0.5 mM isopropyl-β-d-thiogalactopyranoside (IPTG) at 16 °C overnight. The cells were collected and lysed and protein was purified with His-Trap chelating column (GE Healthcare). The purified recombinant proteins were analysed by Coomassie-stained SDS–polyacrylamide gel electrophoresis (PAGE) and desalted into the final storage buffer (100 mM sodium phosphate, pH 7.2) by ÄKTA pure (General Electric, USA). Proteins were concentrated to 2∼4 mg/ml using Ultra centrifugal filters. Aliquots were flash-frozen and stored at -80 °C.

### Cell culture

Mouse neuroblastoma (N2a) cells were cultured in complete medium containing 44.5% Dulbecco’s modified Eagle’s medium (DMEM) 1× with glucose (4.5 g/L), 44.5% Minimum Essential Medium (MEM) Alpha 1×, 10% fetal bovine serum (FBS), and 1% antibiotics (penicillin/streptomycin). Mouse embryonic fibroblast (MEF) cells were cultured in DMEM containing 10% FBS, 1% non-essential amino acid (NEAA) cell culture supplement, 1% antibiotics (penicillin /streptomycin). CHO-K1/CHO-677 cells were cultured in F12K medium containing 10% FBS and 1% antibiotics. All cells were cultured at 37 °C with 5% CO_2_ in a humidified incubator.

### Bioactivity validation of the eGFP-bFGF

MEF (BALB/C-3T3) cell proliferation was measured using Cell Counting Kit-8 (Beyotime Biotechnology, China). Cells were seeded in a 96-well plate (3500 cell/well/100uL) in DMEM supplemented with 10% FBS and allowed to adhere for 24 h at 37°C. The medium was changed with serum-free DMEM containing various concentrations (0, 0.01, 0.1, 0.5, 1, 5, 10 and 100 ng/mL) of either fusion recombinant eGFP-bFGF or commercial bFGF (SonoBiological, China). After 48 hours, the supernatants were removed, and 100 μL of CCK-8 working solution was added to each well for another 1 h at 37 °C. The CCK-8 working solution was prepared with CCK-8 stock solution and DMEM medium at a 1:10 ratio. The optical density (OD) at 450 nm was measured using SpectraMax M2 microplate reader (Molecular Devices). Each group was performed with three replicates.

### Imaging of bFGF phase separation *in vitro*

Purified proteins were diluted to 0.5-1 mg/ml and desalted into assay buffer (8 mM Na_2_HPO_4_, 2 mM KH_2_PO_4_, 136 mM NaCl and 2.6 mM KCl, pH 7.2). For the phase separation of eGFP-bFGF, 5 μM of the protein was mixed with increasing PEG-8000 concentrations (0% – 20% w/v). For eGFP-bFGF-heparin condensate formation, unless specified, 5 μM of eGFP-bFGF were mixed with heparin in the assay buffer. All operations were performed on ice. The mixed protein solution was immediately loaded into a 96-well plate and incubated for indicated time at the indicated temperature before imaging analysis. Images were captured with a Leica SP8 confocal microscopy with a 100× objective (oil immersion).

### Turbidity assay

Proteins were prepared as described above. The protein solution was mixed with various concentrations of heparin (0∼5 µM) and PEG-8000 (0%–20% w/v) in the assay.

All operations were performed on ice. The mixed protein solution was immediately loaded into a 384-well plate and incubated for 5-7 hours at 4 °C before measuring. Turbidity was measured by absorption at 277 nm in 384-well plates using a SpectraMax M2 microplate reader (Molecular Devices). All samples were examined in triplicates (N = 3).

### Analysis of bFGF oligomers by SDS-PAGE

eGFP-bFGF (5μM) was mixed with various concentrations of heparin or de-O-sulphated heparin (GlycoNovo Technologies, China) containing 1 mM H_2_O_2_ or 5 mM DTT on ice. The samples were incubated at 4 °C in a final volume of 100 μL of the assay buffer. After 10 hours, N-ethyl maleimide (100 μM final concentration) was used to block remaining thiols of cysteine residues. The samples (30 ul) were mixed with 10 ul of 4×SDS loading buffer with DTT (reducing) or without DTT (non-reducing) respectively, heated for 5 minutes at 100 °C, and analysed with SDS-PAGE.

### Phase separation of bFGF on cell surface

Cells were plated onto an eight-well Lab-Tek chambered coverglass (Thermo Fisher Scientific) and cultured to around 70%. Before imaging, the medium was discarded, and the cells were washed with PBS twice. Then, the protein solution (500 nM) was applied to the cells. Confocal microscopy was performed with an inverted Leica SP8 microscope, equipped with lasers for 405-nm, 488-nm, 552-nm excitation. Images were acquired using a 100×objective.

### Z-stack for Living Cell 3-D rendering

Three-dimensional reconstruction platform containing Z-stack were imaged with an inverted Leica SP8 microscope. Briefly, images were acquired using the 100× oil immersion lens, a pinhole of 1 AU, 488 nm laser with 10% laser power, followed by setting the starting position and end position of Z-stack, 100∼200 Nr. of Steps or 1μm z-step size was selected. These z-stack images were reconstructed with ImageJ.

### Fluorescence recovery after photobleaching of cell surface and *in vitro* condensates

Samples of bFGF phase separation in solution or on cell surface were examined on an inverted microscope (LSM 780, Carl Zeiss, Germany) equipped with a confocal spinning disk unit (CSU-X1; Yokogawa, Tokyo, Japan) and a Zeiss 100× oil immersion lens. A field (approx. 0.06 μm for the formed droplet in vitro and 0.04 μm for the punctate of cell surface) was bleached for 15 ns with 100 % laser power of a 488-nm or 405 lasers (1 AU) respectively. After being photobleached, images were acquired at a rate of 0.97 s (in solution) per frame or 1.26 s (on cell surface) for 500 s. The fluorescent intensity of bleached area over time was calculated by Zeiss Zen. Signals were normalized with pre-bleached as 100% and 0s after bleach as 0. At least three FRAP curves were averaged to produce ach FRAP curve by Graphpad prism 7.0.

### Heparinase digestion

Cells were treated with heparinase III (0.2 U/mL, GlycoNovo Technologies, China) for 2-4 hours at 37°C. Then the cells were washed with phosphate-buffered saline (PBS) for three times before further treatments.

### Immunofluorescence staining and live-cell imaging

The cells were plated on an eight-well Lab-Tek chambered coverglass at a density 2×10^4^∼5×10^4^ cells/well for N2a and CHO cells and 2000∼4000 for MEF cells in 200 ul medium and cultured for 24 hours, the culture medium was discarded, and cells were washed twice with PBS. For detection of cell surface heparan sulphate, the cells were incubated with an anti-HS antibody (10E4, USBiological, USA) diluted 1:100 in medium containing 1% BSA for 2 hours at 37 °C. After three washing steps with PBS, the cells were incubated with Alexa Fluor 647-conjugated secondary antibody (Jackson Immuno Research, USA). For detection of cell surface FGFR1, the cells were stained with anti-FGFR1 antibody conjugated with Alexa Fluor 405 (Novus Biologicals, USA) for 1 hours at 37 °C. The immune stained cells were examined under a confocal laser scanning microscopy using an inverted Leica SP8 microscope, equipped with lasers for 405-nm, 488-nm, 552nm, 638nm excitation. Images were acquired using a 100×objective.

### Western blot analysis

MEF cells (BALB/C-3T3) were plated on 6-well plates with a density 2×10^5^ cells/well in 2 ml medium and cultured for 48 hours. Then cells were switched into serum-free medium and cultured for 24 hours. Cells were next treated with 200 mIU/mL of Heparinase III (GlycoNovo Technologies, China) for 3 hours. After that, 500 μg/mL of Heparin and 0.1-10 ng/mL of eGFP-bFGF were added into the medium respectively, incubated for 1 hours. Cells were washed three times with PBS before collected and lysed with RIPA lysis buffer (Beyotime Biotechnology, China). The lysates were centrifuged for 10 minutes at 12,000 rpm and the supernatants were used for Western blot. Anti-ERK1/2 antibody (Santa Cruz Biotechnology, USA) and anti-p-ERK 1/2 (pT202/pY204.22A, Santa Cruz Biotechnology, USA) was used for the detection of total ERK and phosphorylated ERK. Horseradish peroxidase–linked anti mouse IgG (Beyotime Biotechnology, China) was used as secondary antibody. The signals were developed using BeyoECL Plus regent (Beyotime Biotechnology, China) and imaged with Chemiscope mini imaging system (CLINX, China). For protein loading control, vinculin antibody (Santa Cruz Biotechnology, USA) was used. The results were analysed with ImageJ.

## Acknowledgements

This work was supported by funding from National Natural Science Foundation of China (91853116, 22077010 and 21672019 to S-Z. L), Natural Science Foundation of Beijing, China (2204088 to S.X.), Young Scientists Fund of the National Natural Science Foundation of China (21907007 to S.X.), the Fundamental Research Funds for the Central Universities (buctrc201920 to S.X., BAIC201814 to J.L.) and the Swedish Research Council (to J.L.). We thank Dr. Yunxiao Zhang, Dr. Shuibing Chen and Dr. Yongxiang Chen for help editing the manuscript.

## Ethics declarations

### Author contributions

S.X. and S.-Z.L. designed the experiments, analysed the data, and wrote the manuscript. F.Z. and T.Z. performed the experiments and analysed the data. H.Z. helped the cell culture and assays. X.W. and L.C. helped preparing the protein. S.-Z.L., S.X., J.L. and T.T. supervised the whole project.

